# Quantification of double stranded DNA breaks and telomere length as proxies for corneal damage and replicative stress in 64 human keratoconus corneas

**DOI:** 10.1101/244871

**Authors:** Robert PL Wisse, Jonas JW Kuiper, Timothy RDJ Radstake, Jasper CA Broen

**Affiliations:** Department of Ophthalmology, University Medical Center Utrecht, The Netherlands; 1: Department of Ophthalmology. 2: Ophthalmo-Immunology group, Laboratory of Translational Immunology, Department of Immunology, University Medical Center Utrecht, The Netherlands; 1: Department of Rheumatology & Clinical Immunology. 2: Laboratory of Translational Immunology, Department of Immunology, University Medical Center Utrecht, The Netherlands

**Keywords:** keratoconus, DNA damage, Alu elements, telomere length, hTERT

## Abstract

**Purpose:** The pathogenesis of keratoconus (KC) is multifactorial and associated with oxidative stress and subsequent DNA damage. The aim of this study was to investigate differences in DNA damage and replicative stress in patients with KC, and in both healthy and diseased controls.

**Methods:** Sixty-four corneal buttons were obtained from 27 patients with KC after corneal transplant surgery, 21 patients with a decompensated graft (DG), and 16 healthy controls (HC). The amount of intact Alu elements per genome copy as measured by qPCR was used to quantify intact DNA. Telomere length was measured as a proxy for replicative stress. In addition, telomerase reverse transcriptase *(hTERT)* gene expression level was assessed.

**Results:** Mean (±SD) DNA damage was similar between the KC (5.56 ±14.08), DG (3.16 ±8.22), and HC (3.51 ±6.66) groups (*P*=0.807). No associations were found between DNA damage and patient age (*P*=0.523), atopic constitution (*P*=0.240), or contact lens wear (*P*=0.393). Telomere length differed (*P*=0.034), most notably in the KC group, and *hTERT* was not detected in any corneal sample. Three cross-linked (CXL) KC corneas did not contain significant more DNA damage (2.6x, *P =* 0.750).

**Conclusions:** Based on these findings, differences in actual corneal DNA damage in KC could not be identified, and the longer telomere length in KC did not support replicative stress as a major etiological factor in the pathogenesis of KC. Future longitudinal investigations on KC etiology should assess progressive early cases to better comprehend the cellular and molecular processes preceding the archetypical morphological changes.

**Precis:** Oxidative stress is allegedly linked with the development of keratoconus. Whether these stressors actually lead to persisting DNA damage and replicative stress is debated. DNA damage was comparable with control samples, and a shortened telomere length was not identified.

## Introduction

Keratoconus (KC) is a corneal condition that can lead to refractive myopia, irregular astigmatism, corneal thinning, and poor visual acuity due to the hallmark “cone-like” shape of the cornea and—in advanced cases—corneal scarring.^1^ Both environmental (e.g., eye-rubbing, atopic constitution, etc.) and genetic factors have been linked to hypersensitive oxidative stress responses at the ocular surface.^2-4^ Currently, crosslinking (CXL; a combination of local ultraviolet (UV)-radiation and riboflavin photosensitizing eye-drops) is regarded an established treatment to prevent disease progression and much research is devoted to study UV-related treatment effects.^5^ However, DNA damage induced by UV radiation has also been suggested as a possible causative factor in the development of KC.^6,7^ UV radiation can damage DNA, leading to breaks in the double stranded DNA, and the ocular surface is heavily exposed to the potentially detrimental effects of UV radiation on DNA integrity.^7,8^ Consequently, the cornea has several robust intrinsic defense systems against UV-induced damage and reactive oxidative species (ROS) in particular. Indeed, studies have found altered activity of several enzymes in the corneas of patients with KC, including the enzymes superoxide dismutase^9^, aldehyde dehydrogenase^10^, catalase^11^, cathepsin^11^, glutathione reductase, transferase, and peroxidases^10^; this altered enzyme activity may therefore contribute to oxidative stress and accumulation of damage.

When DNA damage accumulates, cells usually enter apoptosis. Hence, when there is increased exposure to -for instance-UV light, cells will accumulate DNA breaks rapidly. When DNA repair mechanisms are not able to cope with these breaks, the cells will enter apoptosis, subsequently leading to an increased cell turnover. In keratoconus, one can hypothesize that an increased epithelial apoptosis influences the local homeostasis of the cornea, leading to the loss of stromal collagen fibrils fundamental in the disruption of the normal architecture of the cornea. In addition, increased cellular turnover could be a sign of increased damage to corneal cells, forcing increased cellular replication to replace damaged cells. A well-known measure for the rate of cellular turnover and replicative senescence is telomere length. Telomeres function as a non-coding protective end region of chromosomes. Due to the DNA endreplication inefficiency of polymerases, chromosomes shorten every cell division. After a certain number of divisions, the threshold of attrition is reached, and cells go into apoptosis or enter a senescence state. Telomeres therefore are indicative of the number of divisions the cell lineage has undergone and are a measure of replicative stress. Repair systems exist (i.e. *hTERT*, human telomere reverse transcriptase) and are sometimes upregulated in cells that divide rapidly, such as stem cells.^12^

To answer the question whether corneal cells present in the cornea of keratoconus patients consist of cells refractive to DNA-damage induced apoptosis we quantified the amount of DNA breaks in these cells by measurement of intact *Alu* elements, a proxy for the amount of DNA breaks.^13-17^ The amount of DNA breaks represents a composite measure of the balance between DNA repair and DNA damage. An increase of DNA breaks in keratoconus corneas compared to healthy or decompensated graft corneas could highlight a role for either increased DNA damage, or decreased damage response in this disease.

Secondly, we investigated whether these corneal cells show signs of increased replicative cell turnover by measuring telomere lengths and *hTERT* gene expression as additional parameters of replicative function. Hypothetically, short telomeres in corneal cells could point to a constitutional increased replicative rate, preceding the disrupted morphological structure of a keratoconus cornea.

To answer these questions we assessed DNA breaks, telomere length (TL), and *hTERT* expression in 64 human corneal buttons from 27 patients with KC who underwent corneal transplant surgery, 21 patients with a decompensated graft (diseased controls; DG) not related to KC, and 16 unaffected (healthy) post-mortem donor corneas (HC).

## Material and Methods

### Corneal samples

This study was approved by The Medical Research Ethics Committee of the UMC Utrecht. It reviews research protocols in accordance with the Medical Research Involving Human Subjects Act (WMO). The MREC of the UMC Utrecht is accredited by the Central Committee on Research Involving Human Subjects (CCMO) since november 1999. The MREC of the UMC Utrecht is also member of the Dutch union of MRECs (NVMETC). None of the donors were from a vulnerable population and all donors or next of kin provided written informed consent that was freely given.

Twenty-seven cornea samples were obtained from 27 patients who received a corneal transplant for severe KC and one case of pellucid marginal degeneration (the KC group). A second group of 21 corneal samples was obtained from 21 patients who underwent a re-grafting procedure due to a decompensated corneal graft in which the indication for the primary graft was not KC (the DG group). The corneal buttons in the KC and DG groups were embedded in Tissue-Tek (Sakura Finetek USA, Inc., Torrance, CA) immediately after resection and stored at -80°C.

Corneal samples were also obtained from ten healthy controls (the HC group); these samples were obtained from the Euro Cornea Bank (Beverwijk, the Netherlands) and the Department of Anatomy, University Medical Center Utrecht, Utrecht, the Netherlands. Within 24 hours of death, the corneas were prepared from post-mortem tissue obtained from 16 unrelated donors, each of whom had no documented history of KC, ocular inflammation, or vitreo-retinal disease. All patients provided written informed consent after the explanation of the procedure. Informed consent for the post-mortem donation of ocular tissue was provided under the auspices of the head of the Department of Anatomy, University Medical Center Utrecht, the Netherlands. All tissues were acquired in compliance with Dutch law (*Wet op de lijkbezorging*, Art 18, lid 1/ 18–06–2013) and the institutional guidelines established by the University Medical Center Utrecht.

### Clinical data extraction

Additional data was extracted from the patient records and included both the patient history and the preoperative assessment, which included the results of a slit lamp evaluation, Schirmer’s test, and Scheimpflug corneal tomography (Pentacam HR, Oculus GmbH, Wetzlar, Germany). The data available for the HC group were limited to age, gender, and cause of death. Each KC cornea was graded as clear, mildly hazy, or clouded based on slit lamp biomicroscopy. All decompensated grafts were considered clouded, and all healthy corneas were considered clear.

### Assessment of DNA damage

DNA was isolated from the corneal buttons using TRIzol (Life Technologies, Thermo Fisher Scientific, Grand Island, NY, USA); this approach allows for the isolation of small DNA molecules, which can be lost when using column-based isolation techniques. After dissolving the cornea in TRIzol reagent, the RNA was removed, and the original tubes containing the TRIzol reagent and DNA were stored at -20°C. After isolation, double-stranded DNA was measured using a Qubit 2.0 Fluorometer (Life Technologies, Thermo Fisher Scientific). The number of intact Alu elements in the DNA was measured using qPCR and was used as a proxy for quantifying intact DNA per genome copy, as indicated by the single genome marker 36B4.^13^ DNA damage was assessed in the first 33 consecutive samples, and validated in the subsequent 31 samples. Since baseline characteristics and study outcomes varied only marginally, these were reported as one pooled cohort.

### Measurement of telomere length (TL)

Corneal Telomere Length was measured by Bio-rad cfx-96 real-time qualitative polymerase chain reaction (qPCR) detection system in duplicates in two separate experiments. Briefly, the length of telomeres—long repetitive hexamer (TTAGGG) sequences-can be accurately determined by using a calibration curve based on linear serial dilution of a synthetic 84-mer (14 consecutive TTAGGG sequences) oligonucleotide (Geneworks, Adelaide, Australia) with a predetermined molecular weight per reaction (60×10-12 gr of telomere oligomer or 1.36×109 oligomers). The total number of base-pares in the highest standard can be calculated as ((1.36×109 molecules of oligomer) × (84 oligomer length) = 1.18×108 kilo base-pares). The relative telomere length per sample is extrapolated from serial dilutions of the synthetic standard in each qPCR measurement. Similarly, a synthetic standard was also designed for the single copy house-keeping gene 36B4. Absolute telomere base-pares per genome are quantified by subdividing the total number of telomere base-pares from 36B4 (which has only one copy per gene) following; Telomere length/house-keeping gene = telomere length per genome. ^18^ TL quantification was performed in the first 33 consecutive corneal samples.

### hTERT gene expression

*hTERT* gene expression level was quantified using synthesized cDNA (Biorad iScript kit) from RNA that was extracted from the corneal cells, in the first 33 consecutive cornea samples. Quantstudio qPCR apparatus with TaqMan assay (Applied Biosystems, Thermo Fisher Scientific) were used to perform qPCR under conditions as specified by the manufacturer. To normalize the hTERT-gene expression, the housekeeping GUSB and GAPDH genes were included.

### Statistical analysis

The level of DNA damage (measured using the number of amplified Alu elements) was corrected for input DNA and graphed in a box plot; the mean levels of DNA damage and telomere length were calculated for each study group. Statistical analyses were performed using SPSS 21.0 (IBM, Armonk, NY, USA). Outlier analysis was performed by removing samples with DNA damage that exceeded >3 SD; these values were analyzed separately. Differences in DNA damage, telomere length and *hTERT* expression were tested using the one-way independent ANOVA or Kruskal Wallis test. Multiple comparisons were tested using the post hoc Tukey’s test.

## Results

### Study population

The mean (±SD) ages of the subjects in the KC, DG, and HC groups were 42.8 ± 14.8, 67.6 ± 12.0, and 83.6 ± 8.8 years, respectively. Males were overrepresented in the KC group (63.0%), underrepresented in the DG group (33.3%), and equally distributed in the HC group (56.3%). As expected, concurrent atopic disease was more prevalent in the KC group (in 63.0% of patients) compared to the DG group (9.5%); contact lens wear was also more prevalent in the KC group compared to the DG group (77.8% vs. 57.1%, respectively). The KG and DG eyes were similar with respect to the Schirmer’s test results. Four samples in the DG group, five samples in the HC group and one sample in the KC group did not yield sufficient DNA for analysis, thus reducing the effective sample size from 64 corneas to 54. The characteristics of the four DG patients with non-viable samples did not differ from the mean group (data not shown). The donors of the non-viable HC samples were among the oldest samples (the mean age of these four subjects was 87 years). The mean age of the 33 samples used for telomere length/hTERT assessment did not differ materially from the pooled group (KC 40.8 ± 15.2, DG 66.7 ± 11.9, HC 81,5 ± 9.8 years).

### DNA damage

The mean levels of DNA damage in the pooled KC (n=26), DG (n=17), and HC (n=11) groups were 5.56±14.08, 3.16±8.22, and 3.51±6.66, respectively. Log^10^ normalized values were -0.66±1.63, - 0.59±1.54, and -1.01±1.60 respectively (*P*=0.807; Figure 1).

**Figure 1:**
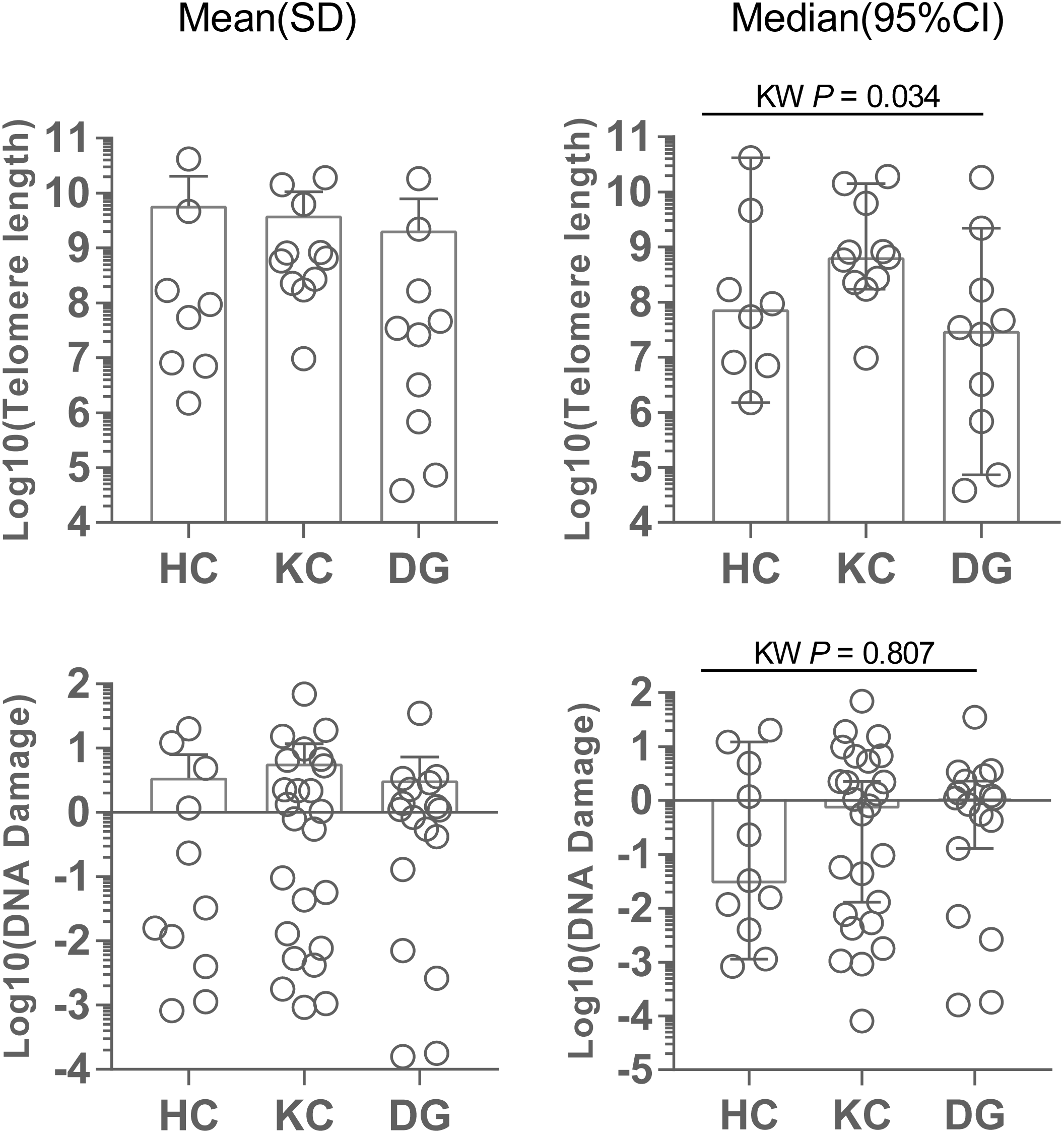
Box plots with overlying scatter plots summarizing the mean and median DNA damage and telomere length (TL) measured in the three study groups. DNA damage was measured as the number of intact *Alu* elements per genome copy. TL is represented as total number of base pairs per genome. Top: Telomere length. Bottom: DNA damage.

The values above represent the total number of breaks in the DNA double strands of all Alu elements per genome copy; thus, considerable variability was observed with respect to the total number of DNA breaks per sample (which ranged from 8.653×10^-5^ to 70.2 in this study). Therefore, we use outlier analysis to normalize the groups; using this approach, we excluded three KC samples, one DG sample, and two HC samples from the analysis (see Table 1). Removing these outliers reduced the average DNA damage in the KC, DG and HC groups to 1.74 ±2.71, 1.19 ±1.24 and 0.71 ±1.61 respectively, but did not affect the mean differences between the groups (*P*=0.407). Similar results were obtained when the results were adjusted for age (data not shown), and we found no significant correlation between age and DNA damage (Spearman’s *ρ*=–0.095, *P*=0.523).

**Table 1:** Summary of DNA damage in the three study groups (N=54*)

|  | N | Mean age | Mean ± SD<br>(all samples) | Multiple<br>comparison<sup>1</sup> | Mean ± SD<br>(outliers removed)<sup>2</sup> | Multiple<br>comparison<sup>1</sup> |
| --- | --- | --- | --- | --- | --- | --- |
| **Keratoconus (KC)** | 26 | 42.7 | 5.56 ± 14.08 | P=0.807 | 1.74 ±2.71 | P=0.407 |
| **Decompensated grafts (DG)** | 17* | 71.4 | 3.16 ± 8.22 |  | 1.19 ± 1.24 |  |
| **Healthy controls (HC)** | 11* | 81.4 | 3.51 ± 6.66 |  | 0.71 ±1.61 |  |
| <sup>1</sup> Independent samples Kruskal-Wallis test. <sup>2</sup> Three outliers were removed from the KC group, one outlier was removed from the DG group, and two outlier was removed from the HC group. * Four samples in the DG group and five samples in the HC group did not yield sufficient DNA for analysis |  |  |  |  |  |  |

Both atopy and contact lens wear are known risk factors for developing keratoconus. In the KC group, seventeen patients had atopy and twenty-one patients wore contact lenses; in the DG group, two patients had atopy and twelve patients wore contact lenses. Therefore, we next investigated the extend of DNA damage among these subgroups. The amount of DNA damage did not differ significantly between the atopic (2.26 ± 2.97) and non-atopic patients (1.26 ± 1.30; P=0.240) or between the patients who wore contact lenses prior to surgery and the patients who did not wear contact lenses (*P*=0.393). We found that the level of DNA damage did not differ significantly from the clear corneas to hazy corneas (*P*=0.418). Clear corneas showed a comparable amount of DNA damage (1.18 ± 2.47) to the mildly hazy corneas (1.80 ± 2.57), and the clouded corneas (1.18 ± 1.16). Three KC cases underwent CXL prior to a grafting procedure. DNA-damage increased two-fold compared to overall outcomes, though this difference was not significant (*P*= 0.750).

### Telomere length and hTERT expression

Telomere length (TL) is represented as total number of base pairs per genome. Four samples did not yield sufficient DNA for analysis. Mean telomere length in KC cases (n=11) was 3.92×10^9^ ± 6.61×10^9^ (log^10^ 8.88 ±0.945), 2.09×10^9^ ± 5.78×10^9^ (log^10^ 7.22 ±1.83) in decompensated grafts (n=10), and 5.83×10^9^ ± 1.46×10^10^ (log^10^ 8.02±1.49) base-pairs in healthy controls (n=8). These differences were statistically significant (*P*= 0.034). Correction for age and gender did not affect these findings. TL was log^10^ normalized to perform these analyses. *hTERT* did not reach detectable levels in any corneal sample.

## Discussion

In this study, we hypothesized that increased DNA damage could be a feature of keratoconus corneal cells, caused either by increased DNA damage or decreased DNA damage responses. In addition, we hypothesized that increased damage could lead to increased replicative senescence or that increased replication of corneal cells on itself could underlie the epithelial remodeling in keratoconus. In this particular large sample, we found that the level of DNA damage and telomere length in the corneas of patients with keratoconus was similar to two control groups (patients with a decompensated graft not related to KC and healthy donor subjects). Telomerase reverse transcriptase *(hTERT)* was not expressed in any corneal sample. In addition, we found no significant correlation between DNA damage, telomere length and either age, gender, atopic constitution, or the use of contact lenses. Telomere length and *hTERT* expression levels have not been assessed previously in corneal tissue.

Several lines of evidence support the notion that keratoconic eyes have altered anti-oxidant function and/or an inadequate DNA repair system.^19^ DNA breaks are a feature of severe DNA damage and can be caused by various agents such as UV light, ROS or chemicals. When these breaks occur, cells initiate DNA repair responses. After this response there are three possible outcomes; the break is repaired, there is too much damage and the cell goes into apoptosis or, as recently discovered, the break persists and the cell goes into a senescent state.^20^ Taking this into mind, the DNA breaks assessed in our study both represent recent non-repaired breaks as well as accumulated unrepaired breaks that did not lead to apoptosis. Following our results, we do not have an indication of increased DNA damage in corneal cells of keratoconus patients compared to healthy and diseased controls.

This does not rule out a contribution to KC development by more specific mechanisms. For example, Atilano et al. reported increased damage to mitochondrial DNA (mtDNA) in the corneas of patients with KC. However in another study, the corneal epithelium of KC patients did not appear to have increased DNA damage.^6,21^

The majority of DNA sampled in our specimens should be regarded of epithelial origin and these short lived mitotic highly active cells are replenished form the stem cell niche, located at the corneal limbus. The limbal region of the cornea is not explanted in routine corneal transplant surgery, though the epithelial cells should represent the genomic make-up of their progenitor stem cells.^22^ This has been well studied and established for telomere attrition; due to the DNA end-replication inefficiency of polymerases, chromosomes shorten every cell division. In line with this, the epithelial cells will reflect the number of cell divisions made by their progenitor cells. Since the differences we found between telomere lengths points towards longer telomeres in KC patients, we assume that there is not a clear indication of increased regeneration of corneal epithelial cells in keratoconus. Naturally, biologically younger samples are expected to have longer telomeres, and statistical correction for age effects was hampered by the poor age distribution among the three sample sets. Since this is the first report on telomere length in corneal tissue, no comparison with younger healthy subjects is available. Mallet et al. demonstrated that corneal epithelial cells have very efficient DNA repair mechanisms, and are less prone to UV-induced apoptosis. Possibly the need to remove highly damaged cells is hereby reduced.^23^ Overall, it seems that the morphological characteristics of keratoconus are not mediated through accumulated DNA damage, aberrant DNA repair systems, or aberrant telomerase systems. Should the peculiar pre-clinical epithelial remodeling rather be attributed to inflammation in the corneal micro-environment?^24,25^

Interestingly, three patients with KC in our study underwent an epithelium-off corneal crosslinking procedure with UV-A irradiation years prior to the grafting procedure, and these patients only showed 2.6x higher level of DNA damage compared with the other samples. Therefore our data do not support a putative link between induced UV-A exposure and DNA damage, which was suggested in a recent report on a case in which crosslinking was associated with intraepithelial neoplasia.^26^

Finally, the current study investigated samples derived from patients that generally suffered from more advanced stages of corneal disease, which are inherently more eligible for corneal transplant surgery. To circumvent this, further investigations should preferably take into account early stages of KC and a longitudinal study design. This could be facilitated by studying corneal epithelium harvested at a CXL procedure, a promising tissue that we are currently exploiting in our laboratory to further explore the dynamics of KC pathology.

## Conclusions

This study is the first to quantify DNA damage using intact *Alu* elements and assess telomere length and *hTERT* expression in corneal tissues. In summary, we found no signs of increased DNA damage or replicative stress in corneal samples obtained from patients with keratoconus compared to healthy controls and patients with decompensated grafts. Thus, the link between DNA repair system dysfunction or accumulated genetic changes due to oxidative stress or UV radiation in the development of progressive KC is unclear. Future longitudinal research should be aimed at younger samples of progressive KC. A thorough analysis of local and systemic auto-inflammatory changes should further elucidate our understanding of KC development.

## Acknowledgments

We thank Rina Wichers, Sanne Hiddingh, Ronald Bleys, and Jan Beekhuis for their assistance in processing the corneal samples.

